# Genetics of adaptation of the ascomycetous fungus *Podospora anserina* to submerged cultivation

**DOI:** 10.1101/488486

**Authors:** Olga A. Kudryavtseva, Ksenia R. Safina, Olga A. Vakhrusheva, Maria D. Logacheva, Aleksey A. Penin, Tatiana V. Neretina, Viktoria N. Moskalenko, Elena S. Glagoleva, Georgii A. Bazykin, Alexey S. Kondrashov

## Abstract

*Podospora anserina* is a model ascomycetous fungus which shows pronounced phenotypic senescence when grown on solid medium but possesses unlimited lifespan under submerged cultivation. In order to study the genetic aspects of adaptation of *P. anserina* to submerged cultivation, we initiated a long-term evolution experiment. In the course of the first four years of the experiment, 125 single-nucleotide substitutions and 23 short indels were fixed in eight independently evolving populations. Six proteins that affect fungal growth and development evolved in more than one population; in particular, the G-protein alpha subunit FadA evolved in seven out of eight experimental populations. Parallel evolution at the level of genes and pathways, an excess of nonsense and missense substitutions, and an elevated conservation of proteins and their sites where the changes occurred suggest that many of the observed allele replacements were adaptive and driven by positive selection.

**Author summary:** Living beings adapt to novel conditions that are far from their original environments in different ways. Studying mechanisms of adaptation is crucial for our understanding of evolution. The object of our interest is a multicellular fungus *Podospora anserina*. This fungus is known for its pronounced senescence and a definite lifespan, but it demonstrates an unlimited lifespan and no signs of senescence when grown under submerged conditions. Soon after transition to submerged cultivation, the rate of growth of *P. anserina* increases and its pigmentation changes. We wanted to find out whether there are any genetic changes that contribute to adaptation of *P. anserina* to these novel conditions and initiated a long-term evolutionary experiment on eight independent populations. Over the first four years of the experiment, 148 mutations were fixed in these populations. Many of these mutations lead to inactivation of the part of the developmental pathway in *P. anserina*, probably reallocating resources to vegetative proliferation in liquid medium. Our observations imply that strong positive selection drives changes in at least some of the affected protein-coding genes.

**Data Availability:** Genome sequence data have been deposited at DDBJ/ENA/GenBank under accessions QHKV00000000 (founder genotype A; version QHKV01000000) and QHKU00000000 (founder genotype B; version QHKU01000000), with the respective BioSample accessions SAMN09270751 and SAMN09270757, under BioProject PRJNA473312. Sequencing data have been deposited at the SRA with accession numbers SRR7233712-SRR7233727, under the same BioProject.

**Funding:** Experimental work and sequencing were supported by the Russian Foundation for Basic Research (grants no. 16-04-01845a and 18-04-01349a). Bioinformatic analysis was supported by the Russian Science Foundation (grant no. 16-14-10173). The funders had no role in study design, data collection and analysis, decision to publish, or preparation of the manuscript.

## Introduction

Over the last several decades, experimental evolution became an essential part of evolutionary biology, providing an opportunity to observe and study changes that occur in controlled populations in the course of many generations. In particular, the *E. coli* long-term evolution experiment led by Richard Lenski since 1988 yielded a number of important results, including that on morphological evolution of cells (1,2), long-term fitness gain under a constant environment (1,3), and ecological specialization (4,5). The existence of twelve replicas made it possible to study changes in genotypes and phenotypes that occurred in parallel in independently evolving bacterial populations (6–8).

Being easily cultured and fast-growing, *E. coli* serves as a convenient model organism for experimental evolution. Still, the range of questions that can be studied on it and on other prokaryotes is limited by their relative simplicity. Eukaryotes such as protozoans or fungi possess more complex gene regulation, morphology and physiology and have also been used in the studies of experimental evolution (9–12).

*Podospora anserina* is a model ascomycete fungus that possesses a definite lifespan with pronounced phenotypic senescence when grown on solid medium (13). In contrast, under conditions of submerged cultivation, *P. anserina* displays unlimited lifespan with no signs of senescence (14,15). Isolates taken from such cultures revert to senescence, indicating that the unlimited lifespan is not associated with genetic changes, instead being a product of epigenetic changes or environmental influences. Still, such isolates demonstrate increased lifespan when plated on solid medium and become sterile, only rarely forming mostly nonfunctional microconidia and protoperithecia (14,15). After some period of submerged cultivation, *P. anserina* cultures acquire higher rates of biomass accumulation and change their pigmentation. The mechanisms that alter *P. anserina* senescence program and affect its phenotype under such conditions are unknown.

In order to study the genetic aspects of adaptation that takes place during the submerged cultivation of *P. anserina*, we started in 2012 a long-term evolutionary experiment. We used two wild strains of *P. anserina* to establish eight independent experimental populations. Here, we analyse the genetic changes that took place in the first four years of this experiment.

## Results

### Allele replacements

We used two founder genotypes, denoted A and B, to establish eight replicate experimental populations (five from A and three from B, referred to as A populations and B populations). Populations were cultivated via serial passages in the standard synthetic medium M2 as described in Materials and Methods. After the short period of initial adaptation to the conditions of the submerged cultivation, all the populations displayed identical phenotypic changes, including changes in colony morphology and pigmentation (15). In order to identify genetic changes acquired by the populations during the course of the experiment, we sequenced both the founder genotypes and each of the populations at four time points (Fig 1).

**Fig 1.**
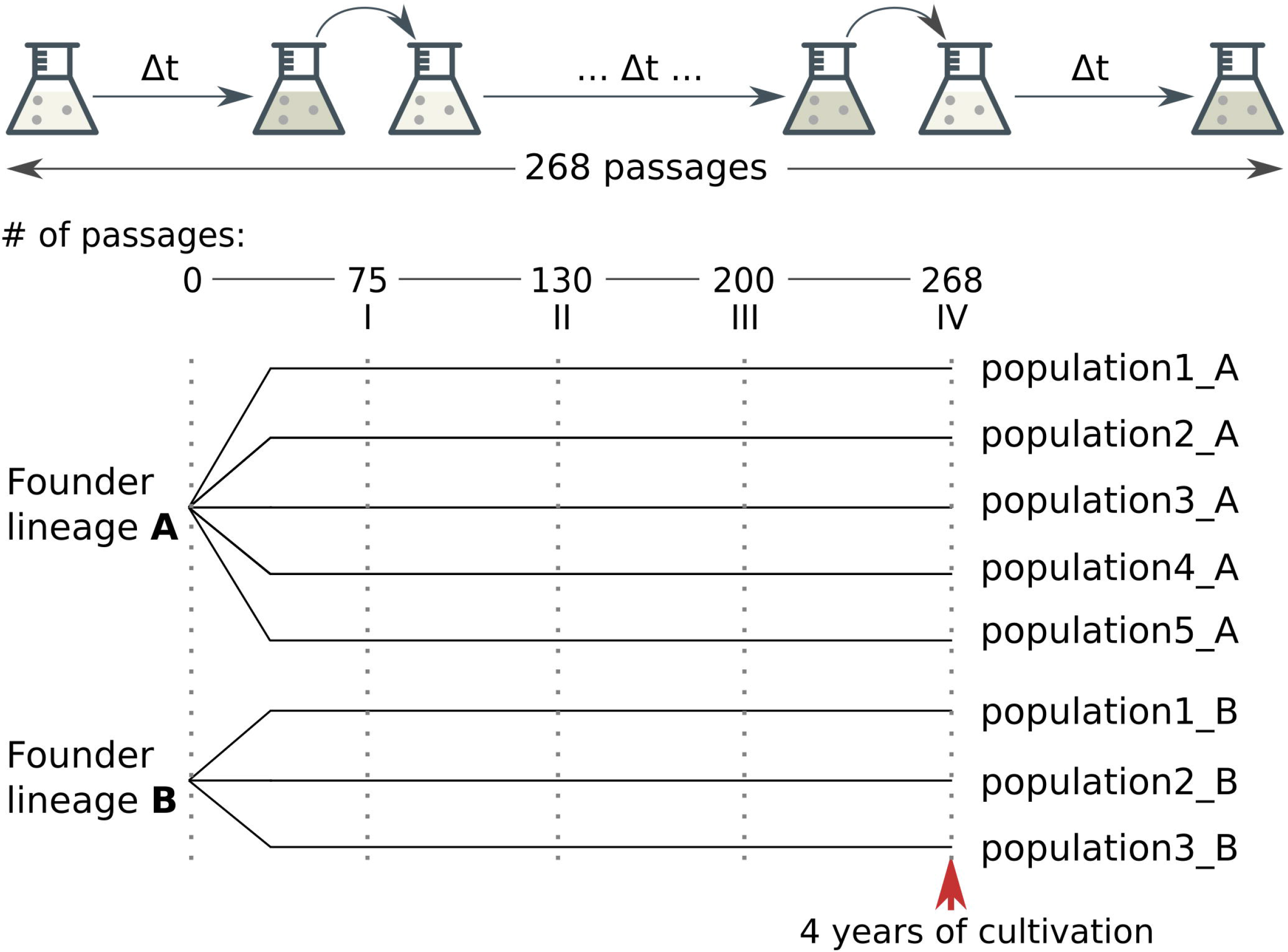
Scheme of the experiment. Eight experimental populations were established from two founder lineages A and B. After 75, 130, 200, and 268 passages, we determined genotypes of the experimental populations (dashed lines; see Methods).

The summary of the observed allele replacements is provided in Table 1 and their full list is presented in Table S1. Our set of allele replacements comprises 85 (73% transitions) and 40 (53% transitions) nucleotide substitutions and 13 and 10 indels in A and B populations, respectively. Among the 148 identified allele replacements, 52 result in an amino acid substitution, 11 are single-nucleotide substitutions leading to a premature inframe stop codon, and 13 cause a frameshift, comprising 35.1%, 7.4% and 8.8% of all allele replacements, respectively.

**Table 1.** Summary of allele replacements that occurred in the experimental populations.

| Type of a replacement | Experimental | Experimental |
| --- | --- | --- |

|  | populations A | populations B |
| --- | --- | --- |
| Total | 98 | 50 |
| By mutation type (single nucleotide mutations): |  |  |
| Transition | 62 | 21 |
| Transversion | 23 | 19 |
| By mutation effect: |  |  |
| Missense nucleotide substitutions | 31 | 21 |
| Synonymous nucleotide substitutions | 6 | 7 |
| Nonsense nucleotide substitutions | 8 | 3 |
| Nucleotide substitutions within noncoding regions | 26 | 5 |
| Nucleotide substitutions within genome regions that cannot be aligned with the reference genome | 9 | 4 |
| Nucleotide substitutions within introns | 3 | 0 |
| Nucleotide substitutions in incomplete proteins | 2 | 0 |
| Short insertions/deletions (including frameshifts) | 13 (9) | 10 (4) |

The dynamics of accumulation of allele replacements is summarized in Fig 2 and Table S2. There are no obvious deviations from their uniform rate in the course of the experiments (linear regression’s time variable p-value is 0.0613). Furthermore, there were no mutators: the numbers of the mutations in each population were all drawn from the same barrel and there were no difference between A and B populations (two-way ANOVA test, see Table S2).

**Fig 2.**
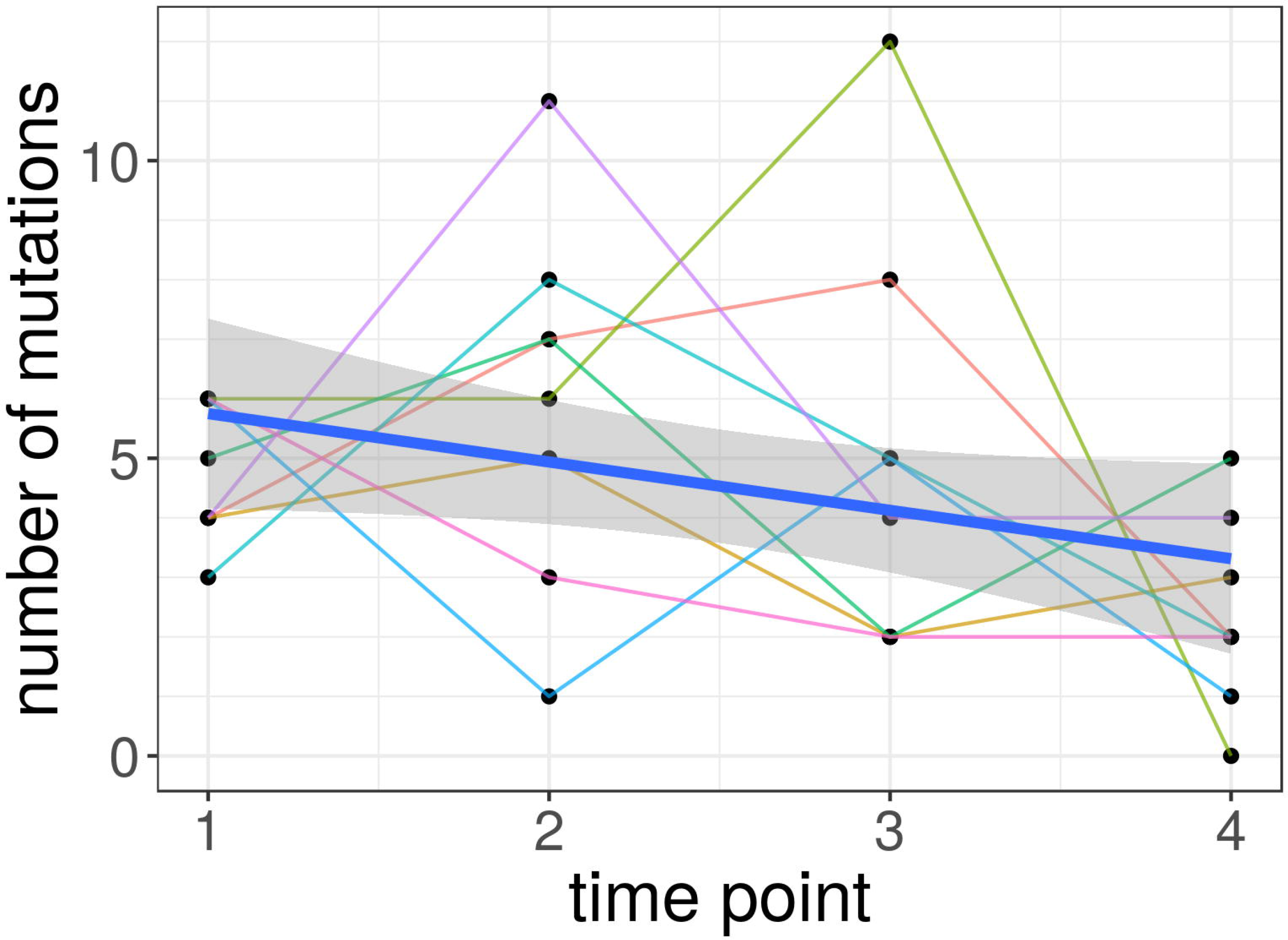
Accumulation of allele replacements during the experiment. Each dot reflects a number of replacements observed in a population at a particular time point. Populations are shown as thin lines. Thick blue line corresponds to the linear regression line with confidence interval depicted as shaded area.

### Distribution of allele replacements within the genome

We then focused on the analysis of single nucleotide substitutions and first computed dN/ dS ratios for populations A and B. dN/dS ratio equals 1.77 and 1.11 for populations A and B, respectively (Table S3). While dN/dS > 1 is commonly thought to indicate positive selection, our estimates may be unreliable because of the scarcity of the data. We next investigated how allele replacements are distributed in the *P. anserina* genome. We produced null-distributions for the number of missense and nonsense substitutions, using permutation procedure (see Materials and Methods). Null-distributions were obtained separately for A and B populations, assuming that mutations strike the genome and become fixed at random, with the uniform per site rates. There are significantly more nonsense substitutions than expected by chance in both A and B populations, and significantly more missense substitutions in B populations (Fig 3).

**Fig 3.**
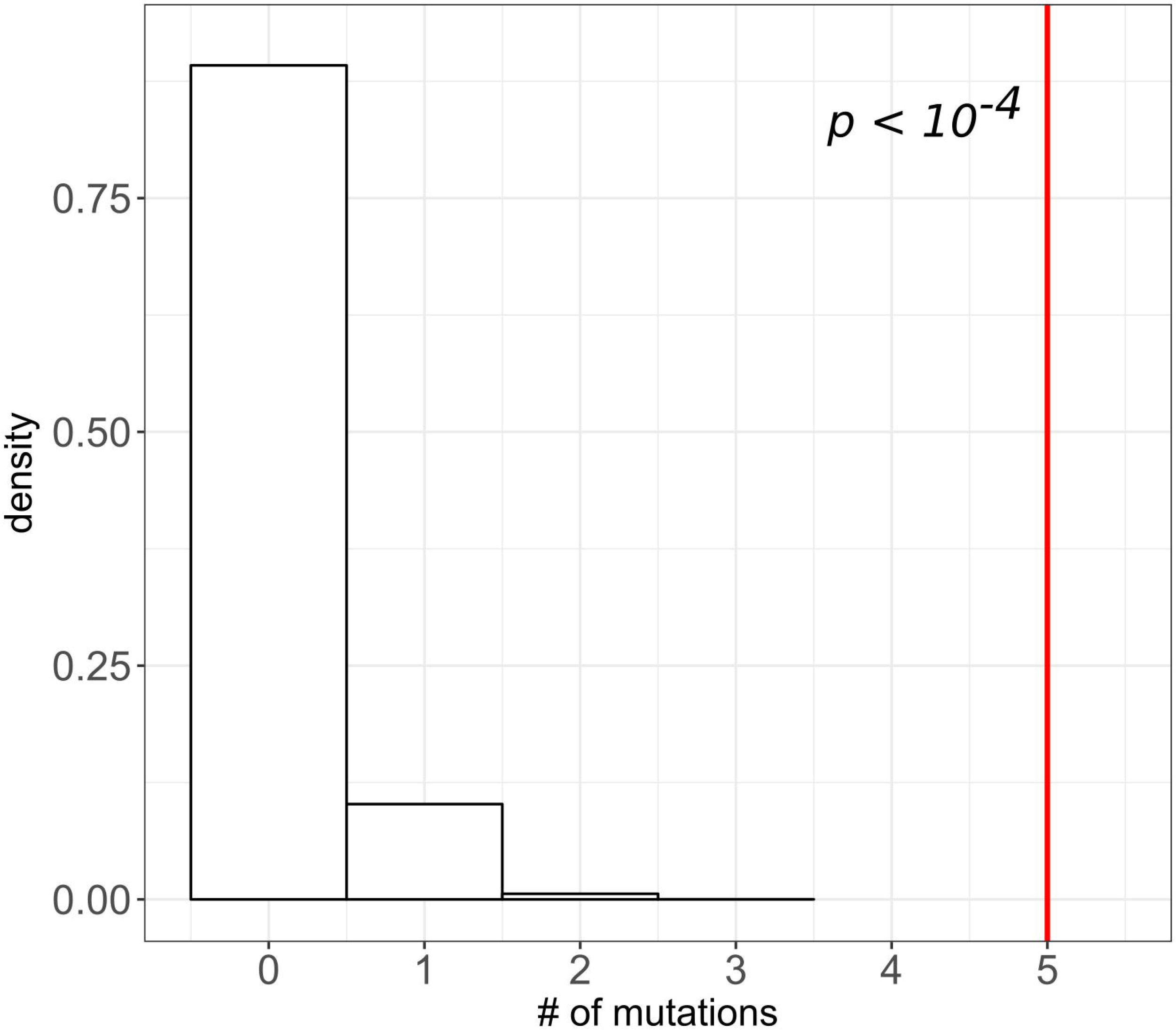
The observed number of missense and nonsense substitutions is greater than expected by chance. Distributions of the expected number of nonsense (A) and missense (B) substitutions in A (left panel) and B (right panel) experimental populations in 10,000 permutations. Red line corresponds to the observed number of nonsense or missense substitutions on (A) and (B), respectively. *p* (p-value) is the area to the right of the red line.

Moreover, despite the fact that there were very few mutations overall, a substantial proportion of protein-altering allele replacements affected the same gene and protein in more than one population. Totally, six proteins were subject to such parallel evolution (Table 2). The probability of observing by chance the same or a higher number of genes carrying just two single-nucleotide protein-altering substitutions by chance is very low (Fig 4), and four out of six proteins that underwent parallel evolution accepted more than two allele replacements. Clearly, positive natural selection was responsible for this phenomenon.

**Table 2.** Proteins affected independently in more than one population.

| Protein identifier and its function | The number of independent allele |
| --- | --- |
|  | replacements |
| Pa_1_10140, Putative transcriptional regulatory protein pro-1 | 2 nonsense substitutions |
| Pa_1_23950, Guanine nucleotide-binding protein alpha subunit encoded by the mod-D gene | 1 nonsense substitutions, 2 missense mutations |
| Pa_3_400, putative protein with unknown function | 6 frameshifts |
| Pa_3_6550, Putative protein homologous to VeA of *Aspergillus nidulans* | 2 missense substitutions |
| Pa_6_3340, Putative developmental regulator flbA | 2 nonsense substitutions, 1 missense substitutions |
| Pa_7_7970, Putative guanine nucleotide-binding protein alpha-1 subunit | 7 missense substitutions |

**Fig 4.**
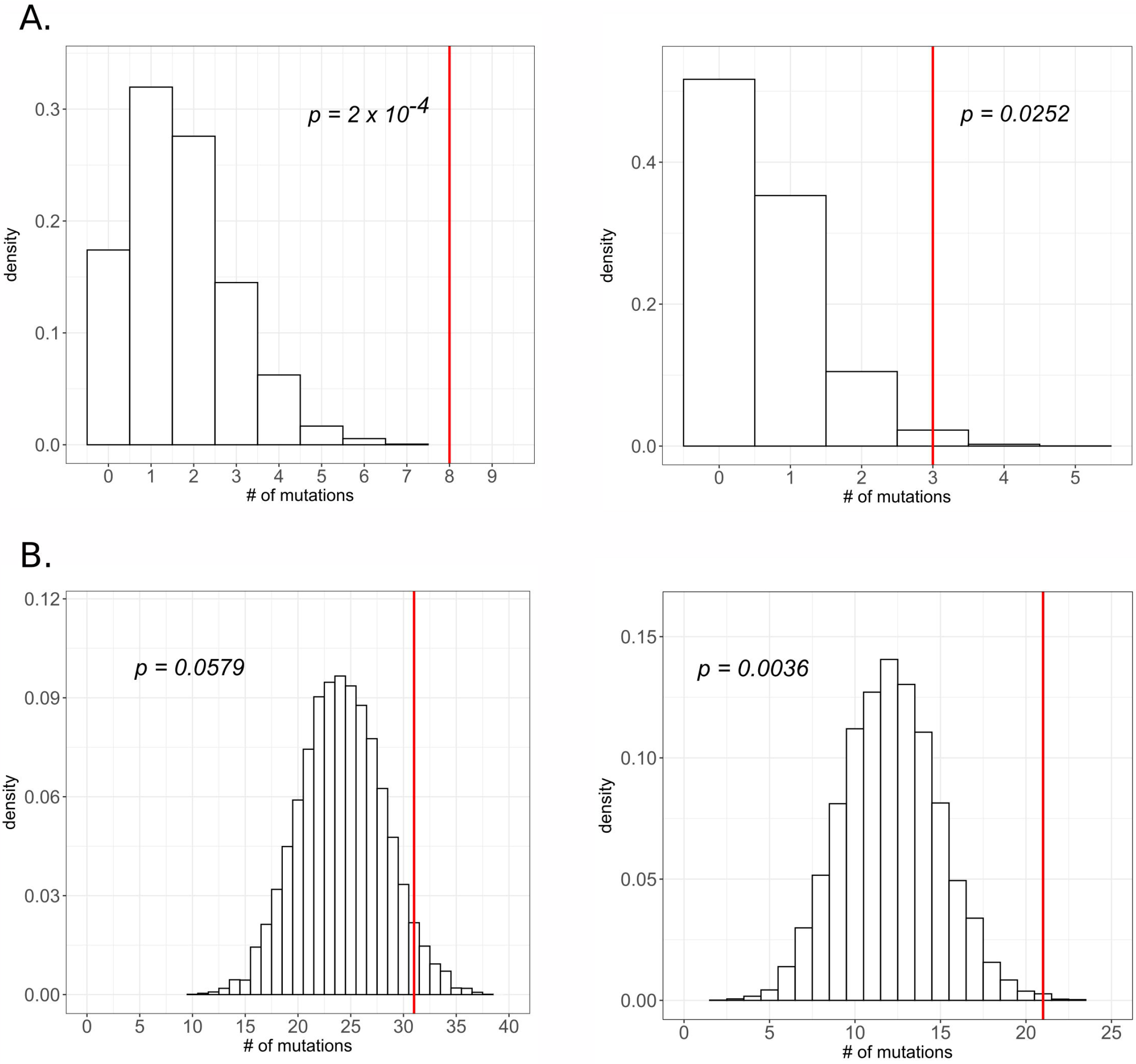
The excess of the number of proteins with parallel changes. Distribution of the number of genes with two protein-altering single-nucleotide substitutions in 10,000 random permutations. Red line shows the observed number of genes with two or more such substitutions occurred.

This parallelism cannot be an artefact of ancestral polymorphism segregating within our lineages, since we conservatively excluded a number of positions with identical parallel mutations in multiple populations founded from the same ancestral lineage. For the same reason, we could lose some actual functional parallelism, making our estimates for the frequency of parallel substitutions conservative. We did observe two identical nucleotide mutations between two lineages formed from two different ancestral lineages (Table S1); those are unlikely to be due to ancestral polymorphism.

### Proteins carrying missense and nonsense substitutions are more conservative

To study the rate of long-term evolution of proteins that evolved in our experiment, we searched for pairs of orthologs pairs encoded by *P. anserina* and *Neurospora crassa* genomes. Out of 10230 proteins in the reference *P. anserina* genome, 4713 have an ortholog in *N. crassa* genome, such that their alignments contain no more than 20% of gaps. Both nonsense and, to a lesser extent, missense allele replacements preferentially occurred in those *P. anserina* proteins that have reliable orthologs in *N. crassa* (Table 3).

Moreover, missense allele replacements within such proteins preferentially occurred at amino acid sites conserved between the two species. While the overall proportion of such sites for 4713 alignments of reliable orthologs is 62%, 87% (27 out of 31) of replaced amino acids occupied conservative sites (p = 0.003).

**Table 3.** Statistics of orthologization of *P. anserina* И *N. crassa* genomes.

|  | Total # of proteins | # of proteins carrying nonsense substitutions | # of proteins carrying missense substitutions |
| --- | --- | --- | --- |
| # of proteins in *P. anserina* genome | 10230 | 9 | 44 |
| # of such proteins that have a reliable ortholog in *N. crassa* | 4713 | 9 (0.000940) | 27 (0.048609) |

Values specified in parentheses correspond to p-values of Fisher's exact test

### Possible phenotypic effects of parallel allele replacements

Proteins that underwent allele replacements in more than one experimental population (Table 2) are of particular interest. Indeed, their evolution must have been adaptive (Fig 4) and, therefore, must have led to changes in the phenotype. Moreover, they probably affected the protein structures radically. Indeed, the differences in physicochemical properties (computed as Miyata’s distance) for pairs of amino acids were higher for missense substitutions that occurred during our experiment than for those that occurred in the course of divergence between *P. anserina* and *N. crassa* mean values 1.979811 vs. 1.472333, two-tail t-test p-value 0.002).

Let us discuss what these changes could be.

The most striking example of parallel evolution is protein Pa_7_7970 which experienced nucleotide substitutions in seven independent populations out of eight. Pa_7_7970 is a putative alpha-1 subunit of a guanine nucleotide-binding protein (Table 2). There are three classes of G protein alpha subunits in ascomycetes fungi (16), each affecting fungal growth and development differently. Pa_7_7970 belongs to class 1 and is 93% identical to its well-studied orthologous protein FadA of *Aspergillus nidulans*. FadA stimulates vegetative growth (17) and probably suppresses sexual reproduction (18), which agrees well with intensive vegetative growth during the experiment suggesting that constitutive activation of Pa_7_7970 may be adaptive under the conditions of submerged cultivation.

Only four sites in Pa_7_7970 evolved in the course of the experiment: arg-178 was substituted in three experimental populations, gly-183 was substituted twice, and ser-206 and lys-210 were each substituted once. All these sites have already been studied. Arg-178 is a catalytic residue crucial for GTP hydrolysis and alpha subunit inactivation (19) and its mutations result in constitutive activation of alpha subunit (20). Gly-183 takes part in the interaction between alpha subunit and regulator of G protein signalling (RGS) that activates GTP hydrolysis, and Gly->Ser substitution (observed in one case) hinders this process drastically (21). Ser-206 and Lys-210 are also involved in the interaction with RGS (22,23) although no specific effects of their mutations have yet been described.

Remarkably, we also observed one missense and two nonsense substitutions in the ortholog of RGS (Pa_6_3340) which belongs to the same pathway as FadA and likely contributes to *P. anserina* adaptation in similar fashion.

Another protein that is worth mentioning is a transcription factor pro-1 (Pa_1_10140) which acquired two loss-of-function allele replacements. pro-1 interacts with promoters of many developmental genes including *esdC* gene that stimulates sexual development and is suppressed by FadA (22,23), and deletion of pro-1 results in sexual sterility. Since *P. anserina* proliferates only vegetatively during the experiment, defects in sexual development pathway may be beneficial.

We observed two missense substitutions in VeA homolog (Pa_3_6550). Although we cannot suggest the effect of two missense substitutions on protein function, VeA in *A. nidulans* positively regulates expression of the previously mentioned *esdC* gene (18) and might affect *P. anserina* metabolism in some way.

We also observed one protein with a surprising amount of parallel evolution via frameshifts. Pa_3_400 protein experienced frameshift in six independent populations. Its homologs in *N. crassa* (24) and *A. nidulans* are involved in asexual reproduction (25). As *P. anserina* lacks asexual reproduction, we cannot suggest the role of Pa_3_400 inactivation in *P. anserina* adaptation, although in *P. anserina* it can be involved in other developmental pathways.

As for Pa_1_23950, it belongs to class 3 alpha subunits of fungal G proteins. Mutations in this protein cause defects in vegetative proliferation in *P. anserina* (26) making it difficult to interpret the observed substitutions in this protein in the context of adaptation.

All proteins discussed above are involved in growth and development processes in various fungi. Taken with the excess of coding substitutions and the number of protein changed in parallel, it favors the possible adaptiveness of some of the observed allele replacements toward the conditions of submerged cultivation. Yet mutant construction is needed to claim it with certainty.

## Discussion

In populations of *P. anserina*, transition to prolonged submerged cultivation inevitably results in the same set of morphophysiological changes (14). These changes always occur early in the course of an experiment (15) and therefore are most likely caused by modifications of epigenetic patterns that alter gene expression. Yet at a longer timescale, one can also expect *P. anserina* to adapt to the conditions that are far from its native environment through fixations of newly acquired mutations.

We analysed small-scale changes that occurred in the *P. anserina* genome in the course of 268 experimental passages. We observed and confirmed the appearance of 125 single nucleotide substitutions and 23 short indels. It is hard to tell which of these mutations are advantageous for submerged cultivation. Experimental populations of *P. anserina* cannot reproduce sexually under conditions of the experiment, and *P. anserina* genus itself lacks the ability to form asexual reproductive structures, macroconidia. Thus, the propagation of our experimental populations occurred exclusively through vegetative proliferation, so that they must possess clonal population structure. Thus, hitch-hiking of neutral and even slightly deleterious mutations that arose in high-fitness genotypes destined to fixation was likely common (27). Nevertheless, two observations clearly indicate that positive selection drove many of the observed allele replacements which, therefore, contributed to adaptation to the experimental environment.

First, evolution of our eight experimental populations often occurred in parallel, with the same gene and protein being affected by a mutation up to seven times (Table 2). Parallel evolution of phenotypes is common in nature, although it does not necessarily involve parallel genetic changes (Steiner et al. 2009). In contrast, in evolutionary experiments parallel genetic changes have being reported several times (28–31). Parallelisms at nucleotide level are rare, and we observed only two such events in Pa_7_7970 gene, which might be mutational hotspots but considering the importance of the mutated protein sites clearly indicate their adaptive role.

Some of the six proteins that underwent parallel changes in our experiment are involved in the same pathways, and all of them seem to be associated with growth and development processes in fungi. A prominent role of nonsense and frameshift alleles, as well as the nature of the fixed missense alleles, indicate that this parallel evolution mostly, if not exclusively, led to inactivation of the affected proteins. At least five missense substitutions in Pa_7_7970, as well as nonsense substitutions in Pa_6_3340, alter developmental pathways probably stimulating vegetative growth and inhibiting sexual reproduction at a very early stage. A similar pattern was observed in experimental yeast populations, where parallel evolution affected genes of the mating pathway and small GTPases of the Ras protein family (29). Loss-of-function mutations of large effects can provide a selective advantage inactivating parts of regulatory/metabolic network that became useless under the current environmental conditions (32). For example, in yeasts mutations resulting in sexual sterility provide advantage in growth rate (33).

Still, the extent of parallel evolution in our experiment is remarkable. Fungi tend to not differentiate in liquid medium (34), so that sexual reproduction was already suppressed before experimental evolution ever begun. Thus, here the situation is very different from parallel loss of eyes in many cave populations of *Astyanax mexicana*, where the cost of maintaining even a useless capacity to see was estimated as up to 15% of resting metabolism (35), which makes its elimination obviously beneficial. In contrast, we observed that positive selection strongly favored mutations that damage a top-level metabolic pathway which already lacks a salient phenotypic expression. Perhaps, before loss-of-function mutations in regulatory proteins occur, some cryptic activity of this pathway remains, and fixation of such mutations allows cells to reallocate their resources to vegetative proliferation. Thus, the role of positive selection in degenerative evolution may be even more prominent than thought previously.

The second reason to believe that many of the observed allele replacements were adaptive is overrepresentation among them of nonsense and missense mutations which must have disproportionally strong impact on function (Fig 3). Moreover, among missense mutations, those that involve radical amino acid replacements and those that occurred at conservative sites and in more conservative proteins are also more common than expected. Because such mutations have a reduced chance of being effectively neutral, this pattern implies that positive selection drove most of their fixations.

Surprisingly, there is still no clarity regarding the role of functionally radical mutations in the adaptive evolution in nature. The radical-conservative ratio of missense substitutions has been studied in the context of both positive and negative selection, and while there are some hints that it correlates with the strength of negative selection (36–38), its relation to positive selection remains ambiguous. Radical-conservative ratio has been proposed as a measure to detect positive selection almost three decades ago (39). However, this measure is vulnerable to various factors not related to selection (40) and studies of different species did not find enough evidences for the excess of radical compared to conservative substitutions at positively selected sites (36,41,42).

A prominent role of mutations at conservative sites and proteins in adaptation to the experimental environment was reported for *E. coli* (43). An obvious possibility is that these, as well as ours, data reflect how adaptation occurs in nature. Indeed, substitutions at conservative sites that occur in the course of natural evolution are enriched with those driven by positive selection (44). Alternatively, a disproportional contribution of radical mutations to experimental evolution may also be due to simplified laboratory environments, under which adaptation mostly involves shedding functions that are no longer needed. Clearly, this issue deserves to be studied further.

At the moment, there is no obvious trend in mutation accumulation rate. Experimental populations acquired two times less mutations between time points 3 and 4 than they did in the previous time interval. However, the reason for that is yet unclear, and further observations might shed some light on this tendency. The experiment lasts for six years already, and to our knowledge, there are not much long-lasting evolution experiments conducted on multicellular organisms. We are planning to continue the experiment and analyse changes in experimental populations of *P. anserina*.

## Material and Methods

### Evolutionary experiment

Two homokaryotic vegetatively incompatible wild strains *s* and *S* (denoted here as A and B) were kindly provided by Annie Sainsard-Chanet and Carole H. Sellem (Département Biologie Cellulaire et Intégrative, Centre de Génétique Moléculaire, CNRS, Gif-sur-Yvette Cedex, France) and gave rise to five and three independent experimental populations, respectively (Fig 1). Experimental populations were maintained via serial passages and were grown in the dark on the standard synthetic medium M2 (15). Submerged cultivation was carried out in 750ml Erlenmeyer flasks with 100 ml of the medium at 27 ± 1°C on a 200rpm rotor shaker. Cultures were transferred to fresh medium every 4-7 days. During the experiment, mycelium samples were plated on agarized (20 g/l) M2 medium in Petri dishes. Initial founder strains are stored on ararized medium.

We denoted 64, 130, 200, and 268 passages as the first, the second, the third, and the fourth time points of the experiment, respectively. Whole-genome sequencing was performed for both the founder populations, for populations B at the points 1, 2, and 4, and for populations A only at the point 4. The allele replacements that were observed from whole genome sequencing data were subsequently verified by Sanger sequencing (see below). Several samples of passage 64 were lost due to contamination; for that reason, we substituted passage 64 with the closest available passage 75 and refer to passage 75 as the first time point in the Results section. The scheme of the experiment is shown in Fig 1.

### Whole-genome sequencing

DNA was extracted using CTAB-based method (45). For library preparation for the first series of samples (founder genotype B and experimental populations B at the first time point), we used Truseq DNA sample preparation kit (Illumina), for the second series of samples (experimental populations B at the second time point) we used NEBNext Ultra DNA kit (New England Biolabs) according to manufacturer instructions. For the third series (founder genotype A and all eight experimental populations at the fourth time point), we constructed PCRfree libraries using Accel-NGS 2S PCR-Free DNA Library Kit (Swift Biosciences). The first and second series were sequenced on HiSeq2000 (Illumina) using Truseq v.3 reagents. The third series was sequenced using Nextseq500 and v.2 reagents.

### De novo genome assembly

Pair-end reads from HiSeq (100bp) and NextSeq (150bp) were trimmed using Trimmomatic (46) with options (ILLUMINACLIP:adapters:2:30:10 LEADING:7 TRAILING7 SLIDINGWINDOW:5:10 MINLEN:50). Duplicate reads were excluded using fastq-mcf (with - D 50 option).

We performed *de novo* genome assembly of both founder genotypes with SPAdes (47) (with -only-assembler option). The initial assemblies were then filtered using the published reference genome (48), v. 6.31, last accessed ~ Dec 2014). We used blastn (49) to find the best hits for each contig in published genome and kept only those contigs that had a local alignment of at least 5kb with published genome with identity of at least 95%. Finally, we performed correction of assemblies using ICORN2 (50). Statistics on assemblies is provided in Table S4.

### Variant calling

For each of the sequenced samples, we mapped its pair-end reads onto the corresponding founder genome using bwa (51) and removed reads that had non-unique alignments. Filtered mappings were used for variant calling. We used samtools (52) to call SNPs and InDels. The resulting calls were filtered by coverage (>=10) and variant frequency (>=80%) to produce high-quality variant set for each of the experimental populations. We then excluded non-unique variants that were supported by at least two reads in more than one experimental population due to extremely low probability of mutations affecting the same base pair in several independent populations. We further masked low complexity regions of founder genomes by repeatmasker (53) and excluded indels that fell into the masked sites and adjacent regions ten base pairs around them. In populations A4 and B2, we observed a couple of mutations in adjacent genomic sites; we excluded these variants as probable dinucleotide mutations.

### Sanger sequencing

Sanger sequencing was used to check the presence of the called variants (138 SNPs and 24 indels) at each of the four time points. Several samples of passage 64 were lost due to *Penicillium* contamination, so we used the closest available passage 75 as the first time point passage. As each of the mutations observed at earlier time points in B populations were always present in later populations, we considered a mutation as verified if it was confirmed by Sanger sequencing in any of four time points (assuming that the mutation that appeared once became fixed at later time points). For B populations, we believed in NGS data for passages 64 and 130 if the mutation was identified by Sanger sequencing at any time point. For A populations, we had to check each of the four time points.

Primers for Sanger sequencing were designed using primer3 (54,55) and tntblast (56) software and synthesized in Evrogen. PCR was performed using Encyclo kit (Evrogen). Amplification products were sequenced in both directions on ABI Prism 3500 sequencer (Applied Biosystems). Chromatograms were validated manually in CodonCode Aligner (57).

The detailed results of Sanger sequencing are included in Table S1. The number of confirmed mutations is provided in Table S2. We were unable to confirm neither the presence nor the absence of 15 mutations and excluded them from any further analyses. The majority of mutations that we failed to validate comprise mutations in noncoding/not annotated regions. Also, there were three mutations in A populations for which we couldn’t track the time of their appearance.

### Variant annotation

We mapped reference CDS sequences (48) (v. 6.31, last accessed ~ Dec 2014) onto the reference genome and obtained coding regions coordinates using Splign (58). Out of 10,635 reference protein-coding genes, we were able to identify 10230 genes in the reference genome that coded complete proteins with proper start and stop codons. We then performed pairwise genome alignment of A and B founder genotype assemblies with published genome using Multiz (59) and identified coding regions coordinates in the assembled genomes using Splign annotation results. 9732 and 9732 (numbers are indeed the same but protein sets differ) complete genes were identified in A and B founder genotypes, respectively, with a tiny amount of incomplete genes (627 and 530 genes for A and B populations, respectively). Based on the produced annotations, the identified genomic variants were classified as exonic, intronic or intergenic. Exonic variants in complete genes (i.e., genes with all of their exons completely identified in the founder genome) were further classified as missense, nonsense or silent mutations in case of SNPs and frameshift or amino acid insertion/deletion mutations in case of indels.

### Permutations

In order to estimate the probability to obtain the observed number of protein-altering variants by chance, we generated null distributions for the number of missense and nonsense mutations in A and B founder genomes using permutations. A permutation procedure was performed as follows. 85 and 40 observed SNP mutations were thrown at random onto A and B genomes, respectively, and annotated as described above. To control for possible confounding effects of mutation context, we added three-nucleotide sequence contexts to permutations. The procedure was repeated 10,000 times and produced null distributions for the expected number of nonsense and missense mutations in each of the founder genomes (Fig 3). To obtain the expected number of proteins evolving in parallel, we combined permutation results from A and B populations and counted the number of proteins that acquired at least two single-nucleotide protein-changing mutations in eight populations during the permutation procedure.

### Protein conservation analysis

We used OrthoMCL (60) to identify orthologous groups between protein sets from *P. anserina* and *N. crassa* (61) reference genomes. Orthologous protein pairs were then aligned using MAFFT (62) (with -localpair option), and for each protein carrying ortholog in *N. crassa* genome, we calculated its conservation as a fraction of conserved positions in the alignment. We also measured the conservation of mutated positions as a fraction of conserved positions among the whole set of mutated positions.

## Supporting information

## Supplementary Information

**Table S1. The summary of the observed allele replacements.** Sheets A1-A5 and B1-B3 contain the results of the Sanger sequencing validation procedure. The two last sheets comprise annotation of the allele replacements in A and B populations.

**Table S2. Accumulation of allele replacements.** Each entry provides the number of newly observed replacements, relatively to the previous time point.

**Table S3. dN/dS calculation.**

**Table S4. Statistics on founder genotypes assemblies.**

